# Theoretical basis for stabilizing messenger RNA through secondary structure design

**DOI:** 10.1101/2020.08.22.262931

**Authors:** Hannah K. Wayment-Steele, Do Soon Kim, Christian A. Choe, John J. Nicol, Roger Wellington-Oguri, Andrew M. Watkins, R. Andres Parra Sperberg, Po-Ssu Huang, Eterna Participants, Rhiju Das

**Affiliations:** Department of Chemistry, Stanford University, Stanford, CA, 94305; Eterna Massive Open Laboratory. Consortium authors listed in Table S1; Department of Chemical and Biological Engineering, Northwestern University, Evanston, IL, 60208; Department of Bioengineering, Stanford University, Stanford, CA, 94305; Department of Biochemistry, Stanford University, Stanford, CA, 94305; Department of Physics, Stanford University, Stanford, CA, 94305

**Keywords:** mRNA therapeutics, mRNA vaccines, RNA structure, RNA degradation

## Abstract

RNA hydrolysis presents problems in manufacturing, long-term storage, world-wide delivery, and in vivo stability of messenger RNA (mRNA)-based vaccines and therapeutics. A largely unexplored strategy to reduce mRNA hydrolysis is to redesign RNAs to form double-stranded regions, which are protected from in-line cleavage and enzymatic degradation, while coding for the same proteins. The amount of stabilization that this strategy can deliver and the most effective algorithmic approach to achieve stabilization remain poorly understood. Here, we present simple calculations for estimating RNA stability against hydrolysis, and a model that links the average unpaired probability of an mRNA, or AUP, to its overall hydrolysis rate. To characterize the stabilization achievable through structure design, we compare AUP optimization by conventional mRNA design methods to results from more computationally sophisticated algorithms and crowdsourcing through the OpenVaccine challenge on the Eterna platform. These computational tests were carried out on both model mRNAs and COVID-19 mRNA vaccine candidates. We find that rational design on Eterna and the more sophisticated algorithms lead to constructs with low AUP, which we term ‘superfolder’ mRNAs. These designs exhibit wide diversity of sequence and structure features that may be desirable for translation, biophysical size, and immunogenicity, and their folding is robust to temperature, choice of flanking untranslated regions, and changes in target protein sequence, as illustrated by rapid redesign of superfolder mRNAs for B.1.351, P.1, and B.1.1.7 variants of the prefusion-stabilized SARS-CoV-2 spike protein. Increases in in vitro mRNA half-life by at least two-fold appear immediately achievable.

**Significance statement:** Messenger RNA (mRNA) medicines that encode and promote translation of a target protein have shown promising use as vaccines in the current SARS-CoV-2 pandemic as well as infectious diseases due to their speed of design and manufacturing. However, these molecules are intrinsically prone to hydrolysis, leading to poor stability in aqueous buffer and major challenges in distribution. Here, we present a principled biophysical model for predicting RNA degradation, and demonstrate that the stability of any mRNA can be increased at least two-fold over conventional design techniques. Furthermore, the predicted stabilization is robust to post-design modifications. This conceptual framework and accompanying algorithm can be immediately deployed to guide re-design of mRNA vaccines and therapeutics to increase in vitro stability.

## Introduction

Messenger RNA (mRNA) molecules have shown promise as vaccine candidates in the current COVID-19 pandemic (1–3) and may enable a large number of new therapeutic applications (4–6). However, a major limitation of mRNA technologies is the inherent chemical instability of RNA. mRNA manufacturing yields are reduced by degradation during in vitro transcription; mRNA vaccines stored in solution require in vitro stability, ideally over months under refrigeration (7); RNA vaccines deployed in developing regions would benefit from increased in vitro stability against high temperatures (8); and after being administered, mRNA vaccines require stabilization against hydrolysis and enzymatic degradation to sustain translation and immunogenicity in the human body (9).

RNA degradation depends on how prone the molecule is to in-line hydrolytic cleavage and attack by nucleases, oxidizers, and chemical modifiers in the RNA’s environment (10–13). Amongst these degradation processes, in-line hydrolytic cleavage is a universal mechanism intrinsic to RNA. Cleavage of an RNA backbone phosphodiester bond is initiated by deprotonation of the 2’-hydroxyl group of the ribose moiety (14) (Figure 1A). The deprotonated hydroxyl group attacks the phosphate to form a pentacoordinate transition state. The formation of this transition state relies on the RNA backbone being able to adopt a conformation where the 2’- hydroxyl group is in line with the leaving 5’ oxyanion. The 5’ oxyanion then departs, leaving behind a 2’,3’-cyclic phosphate and a strand break in the RNA. The same mechanism underlies the action of self-cleaving ribozymes and protein-based nucleases, allowing this conformation to be characterized experimentally and visualized in crystal structures (Figure 1B, structure from ref. (15)).

**Figure 1.**
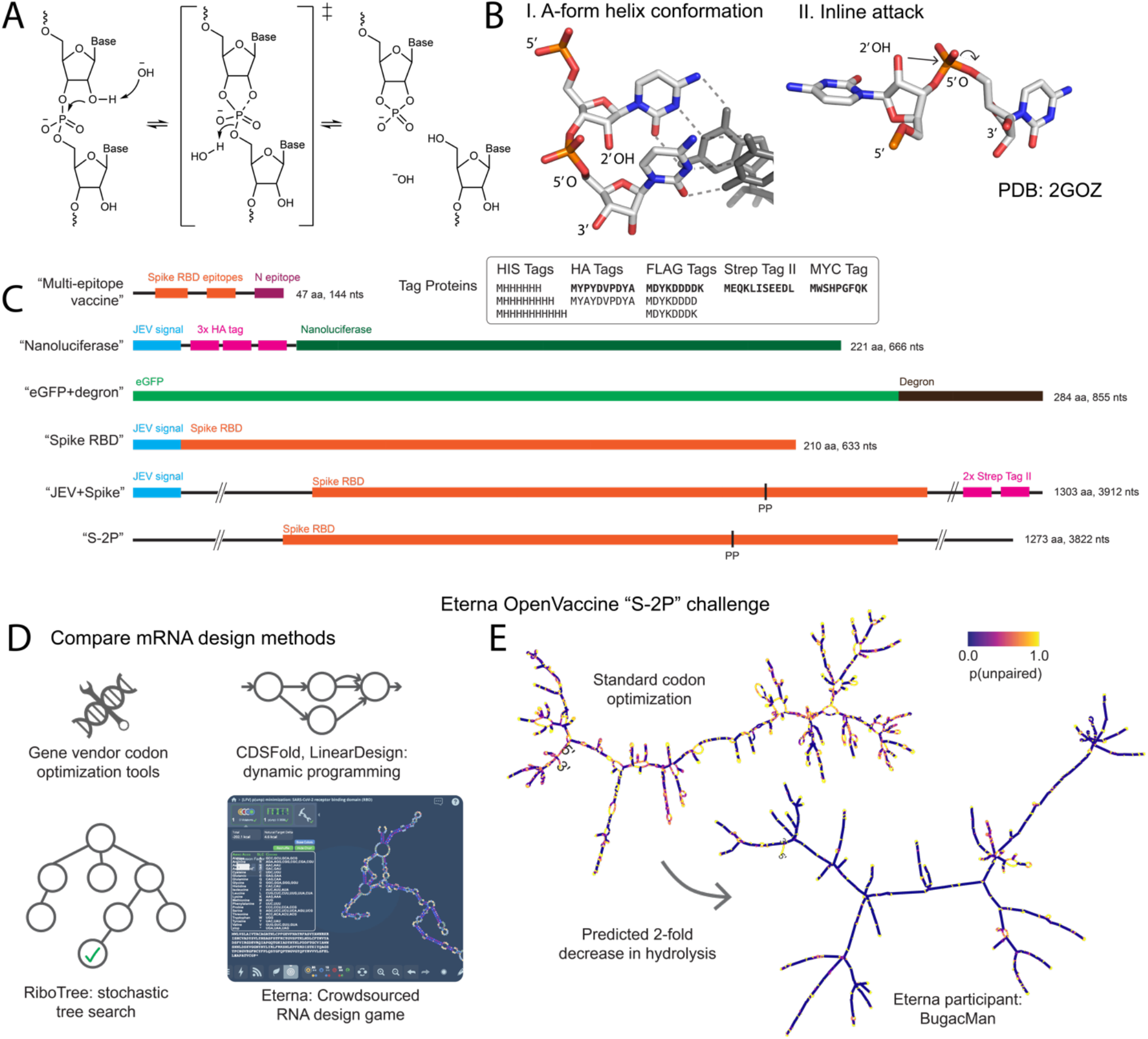
(A) Hydrolysis of the phosphodiester bond in the RNA backbone bond. This mechanism proceeds via an “inline attack” backbone conformation, depicted in (B): the attacking 2’-hydroxyl group is in line with the phosphate group and the leaving 5’ oxygen. (C) Sequence schematics of all mRNA design challenges in this work. (D) We compared the maximum degree of stabilization achievable through a variety of current and novel design methods. (E) mRNAs designed by conventional means for therapeutics are prone to hydrolysis in regions that have high probability of being unpaired (shown in yellow, left panel). A design for an mRNA vaccine encoding the prefusion-stabilized SARS-CoV-2 full spike protein (S-2P) dramatically reduces the probability of being unpaired throughout the molecule (purple, right panel).

Hydrolysis sets a fundamental limit on the stability of mRNA medicines and technologies. The World Health Organization’s target product profile for vaccines calls for formulations that remain effective after a month under refrigeration (2-8 °C) (8). Deployment of mRNA vaccines for infectious disease outbreaks like the current COVID-19 pandemic would benefit from taking advantage of existing supply networks for conventional attenuated vector vaccines, which are set up for pre-filled syringes in saline buffer at near-neutral pH under refrigeration (8). However, model calculations of RNA hydrolysis as a function of pH, temperature, and ionic concentration (16), highlight potential problems for using the same supply networks for mRNA vaccines. Under refrigerated transport conditions (‘cold-chain’, 5 °C, phosphate-buffered saline, pH 7.4, no Mg^2+^) (8), a naked RNA molecule encoding a SARS-CoV-2 spike glycoprotein, with a length of roughly 4000 nucleotides in bulk solution would have a half-life of 900 days, with 98% intact after 30 days, fitting the target product profile for vaccines from the World Health Organization. However, a temperature excursion to 37 °C is predicted to lead to a half-life reduced to 5 days, well under a month. Even if temperature can be maintained at 5 °C, RNAs encapsulated in lipid formulations may be subject to increased hydrolysis if the lipid’s cationic headgroups lower the pKa of the ribose 2-hydroxyl group (17, 18). If pKa shifts as small as 2 units occur, the predicted half-life reduces from 900 days to 10 days, again well under a month (Table 1). Beyond the above considerations for a ~4000 nt mRNA, the longer lengths of RNA molecules (>12,000 nt RNA) being considered for low-material-cost ‘self-amplifying’ mRNA (SAM) vaccines (3, 19) are expected to exacerbate inline hydrolysis. In all conditions described above, the half-life will be reduced by a further 3-fold compared to a non-SAM mRNA. As an example, if during storage or shipment at pH 7.4, the SAM vaccine of length 12,000 nts is subject to an excursion of temperature to 37 °C for 2 days, the fraction of functional, full-length mRNA remaining after that excursion will drop to less than half of the starting RNA (Table 1). Beyond these calculations under storage and shipping conditions, an mRNA vaccine is expected to be highly unstable during in vitro transcription and upon delivery into the human body (half-lives reduced to hours due to presence of Mg^2+^ and physiological temperatures; Table 1). For these reasons, it is desirable to explore principles by which mRNA molecules might be designed to improve their stability against hydrolysis.

**Table 1.** Estimates for RNA degradation using the quantitative model presented by Li & Breaker (14) and the model for AUP presented in this work.

| Simulated condition (0.14 M $[K^+]$) | T (°C) | pH | $[Mg^{2+}]$ (mM) | RNA length (nucleotides) | AUP<sup>a</sup> | Cleavage rate per molecule ($k_{deg}$) ($10^{-7} \text{ min}^{-1}$) | Half-life<sup>b</sup> (days) |
| --- | --- | --- | --- | --- | --- | --- | --- |
| Refrigerated supply chain ('cold chain') | 5 | 7.4 | 0 | 4,000 | 0.4 | 5.1 | 941 |
| Refrigerated supply chain, increased length (SAM RNA) | 5 | 7.4 | 0 | 12,000 | 0.4 | 15.3 | 314 |
| Refrigerated supply chain, pKa shifted by cationic formulation | 5 | 9.4<sup>c</sup> | 0 | 4,000 | 0.4 | 470 | 10.2 |
| Temperature excursion | 37 | 7.4 | 0 | 4,000 | 0.4 | 890 | 5.4 |
| Manufacturing (in vitro transcription)<sup>d</sup> | 37 | 7.6 | 14 | 4,000 | 0.4 | 57,000 | 0.084 |
| Physiological | 37 | 7.4 | 1<sup>e</sup> | 4,000 | 0.4 | 2,000 | 2.4 |

<sup>a</sup> Typical average unpaired probability (AUP) of 0.4 estimated from conventional design methods studied in this work.
<sup>b</sup> Calculated as $t_{1/2} = \ln 2 / k_{deg}$ .
<sup>c</sup> Apparent pH at 2' hydroxyl, simulating pK<sub>a</sub> shift of 2 units induced by complexation with cationic lipid.
<sup>d</sup> Ref. (20), with pH 7.9 of Tris-HCl buffer corrected from 25 °C to 37 °C.
<sup>e</sup> Ref. (21).

One largely unexplored design method to reduce RNA hydrolysis that is largely independent of mRNA manufacturing, formulation, storage, and *in vivo* conditions is to increase the degree of secondary structure present in the RNA molecule. Hydrolysis has been found to be mitigated by the presence of secondary structure, which restricts the possible conformations the backbone can take and reduces the propensity to form conformations prone to in-line attack (22). Indeed, the technique of inline probing takes advantage of the suppression of in-line hydrolysis within double-stranded or otherwise structured regions to map RNA structure (23).

Here, we report a theoretical framework and computational results indicating that structure-aware design should enable immediate and significant COVID-19 mRNA vaccine stabilization. We present a principled model that links an RNA molecule’s overall hydrolysis rate to base-pairing probabilities, which are readily calculated in widely used modeling packages (24–27). We define two metrics: the Summed Unpaired Probability of a molecule, or SUP, and the Average Unpaired Probability, or AUP, which is the SUP normalized by sequence length. By comparing a variety of mRNA design methods (Figure 1D), we provide evidence that both crowdsourced rational design on the Eterna platform and several optimization algorithms are able to minimize AUP for the CDS across a range of mRNA applications. The calculations predict that structure-optimizing designs can achieve at least twofold increases in estimated mRNA half-life compared to conventional design methods (Figure 1E), independent of the mRNA length. Our results furthermore suggest that optimizing for mRNA half-life can be carried out while retaining other desirable sequence or structure properties of the mRNA, such as codon optimality, short stem lengths, and compactness measures, which may modulate in vivo mRNA translation and immune response. The predicted structures of these molecules are robust to a wide variety of perturbations, including temperature excursions, addition of flanking untranslated regions, and nonsynonymous changes in coding sequence, leading us to term them ‘superfolder’ mRNAs.

## Results

### A biophysical model for RNA degradation

Previous studies have explored the design of mRNA molecules with increased secondary structure (28–30), as evaluated by the predicted folding free energy of the mRNA’s most stable structure, but it is unclear if this metric is the correct one when improving stability of an RNA against degradation. Our computational studies are based on a principled model of RNA degradation that suggests an alternative metric. To describe the degradation rate of an RNA molecule, we imagine that each nucleotide at position *i* has a rate of degradation *k_cleavage_(i)*. To focus our presentation, we imagine that the degradation is due to inline hydrolysis, but the framework below generalizes to any degradation process that is suppressed by formation of RNA structure, including digestion by endonucleases for *in vivo* applications. The probability that the nucleotide backbone remains intact at nucleotide *i* after time *t* is

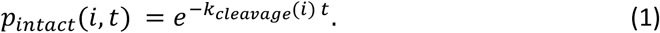

The probability that the overall RNA with length *N* remains intact with no chain breaks after time *t* is then

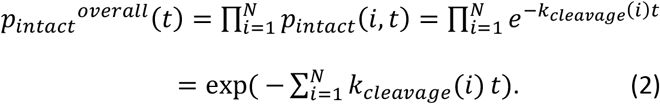

Here, we have assumed that the probability of cleavage at any given position is independent of cleavage events at other positions in the RNA. If this is not true, the expression will still remain correct at times when there are 0 or 1 cleavage events, and that is the time range most relevant for improving RNA stability. Given that assumption, eq. (2) gives an exactly exponential dependence of the overall degradation of the RNA with respect to time *t*:

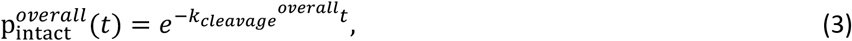

with

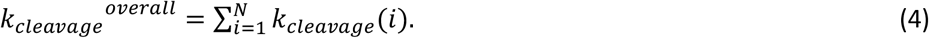

The degradation half-life of the RNA is t_½_ = ln 2 */k_cleavage_^Overall^*.

The rate at which an RNA is hydrolyzed at a specific location along its backbone *k_cleavage_(i)* depends on the ability of the phosphodiester bond to adopt the in-line attack conformation (Figure 1B), or more generally for the RNA to adopt a conformation that can be accessed by a degrading agent, like a protein nuclease. Here and below, we work out the consequences of a simple model reflecting the knowledge that in-line hydrolysis generally occurs at nucleotides that are unpaired but is strongly suppressed by pairing of the nucleotide into double-stranded segments of the secondary structure (22, 23). Since RNA chains fluctuate between multiple secondary structures with a characteristic timescale of milliseconds (31), faster than the degradation rates discussed above, we write the overall cleavage rate as averaged over the equilibrated structural ensemble of the RNA,

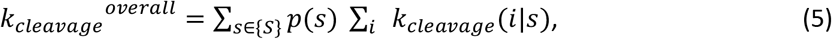

where {*S*} is the full set of structures that the RNA molecule is capable of adopting, and *p(s)* is the probability of forming a structure *s*. The rate of cleavage *k_cleavage_(i|s)* at each position *i* within a structure *s* will, in general have a complex dependence on the sequence and structural context. For example, the cleavage rate will depend on whether the nucleotide is in a hairpin loop, where it is in within the loop, whether other loop nucleotides might promote in-line cleavage through acid-base catalysis, whether the loop has non-canonical pairs, etc. Without additional empirical knowledge, we assume that the cleavage rate for unpaired nucleotides can be approximated by a constant rate *k_cleavage_^Unpaired^* if nucleotide *i* is unpaired, and zero if paired. Then eq. (5) becomes:

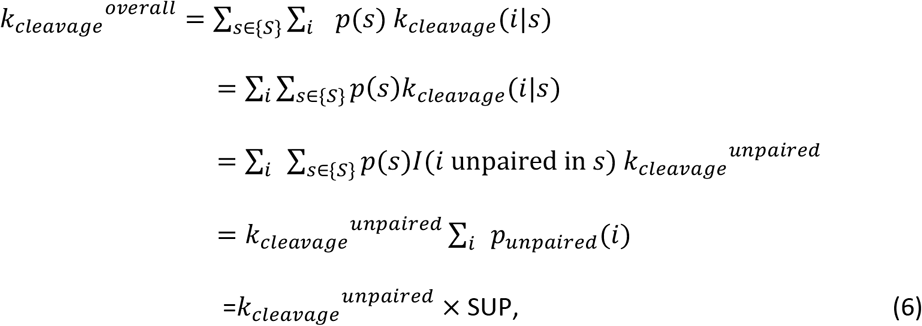

where *I(i unpaired in s)* is 1 if nucleotide *i* is unpaired in the structure *s* and 0 otherwise. In the last line of (6), we introduce the definition of Sum of Unpaired Probabilities (SUP),

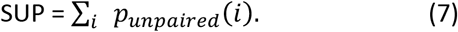

Overall, the total rate of cleavage may be approximated as this measure, the sum of unpaired probabilities across all nucleotides of the RNA, multiplied by a constant *k_cleavage_^unpaired^* that reflects the average cleavage rate of an unpaired nucleotide.

It is important to point out that the total rate scales with the sum of the unpaired probabilities of the RNA’s nucleotides – longer RNA molecules are expected to degrade faster in proportion to their length. This relation is better reflected by a rearrangement of (6) to:

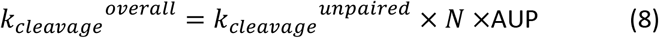

where the average unpaired probability (AUP) is

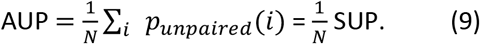

The AUP value is a number between 0 and 1 that reflects the overall ‘unstructuredness’ of the RNA. Lower values correspond to lower probability of being unpaired, and therefore RNA molecules less susceptible to degradation.

In these last expressions (eqns. 6-9), *p_unpaired_(i)* can be predicted in most widely-used RNA secondary structure prediction packages, which output base pair probabilities *p(i: j)*, the probability that bases *i* and *j* are paired. Then the probability that any nucleotide *i* is unpaired in the RNA is given as

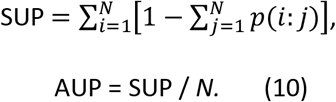

Under this model, it becomes possible to computationally study the question of how much an RNA might be stabilized if it is redesigned to form stable secondary structures, which we describe next.

### Small mRNA models reveal discrepancy in sequences optimized for SUP vs. sequences optimized for codon optimality or minimum folding free energy

To investigate the possible dynamic range in degradation lifetimes for mRNA, we started with mRNA design problems that were small enough to be tractable, i.e., all mRNA sequences that code for the target amino acid sequence could be directly enumerated and studied. We selected a collection of short peptide tags that are commonly appended or prepended to proteins to enable purification or imaging: His tags of varying lengths, human influenza hemagglutinin (HA) tag, Strep-tag II, FLAG fusion tag, and Myc tag sequences (32). We enumerated all the mRNA sequences that encode each protein. For reference, we also generated sequences from online codon optimization tools from gene vendors IDT, GENEWIZ, and Twist (see Methods). As expected, the predicted degradation rates of solutions from these vendor sequences, which do not take into account mRNA in vitro stability, were higher than the minmal degradation rate computed for all enumerated sequences (Figure 2A, Figure S1). We calculated the change in AUP obtained between the average of all the vendor-returned sequences and the minimum AUP solution for tag proteins, and found fold-changes in AUP ranging from 1.27 to 2.23-fold decrease in AUP (Table 2).

**Figure 2.**
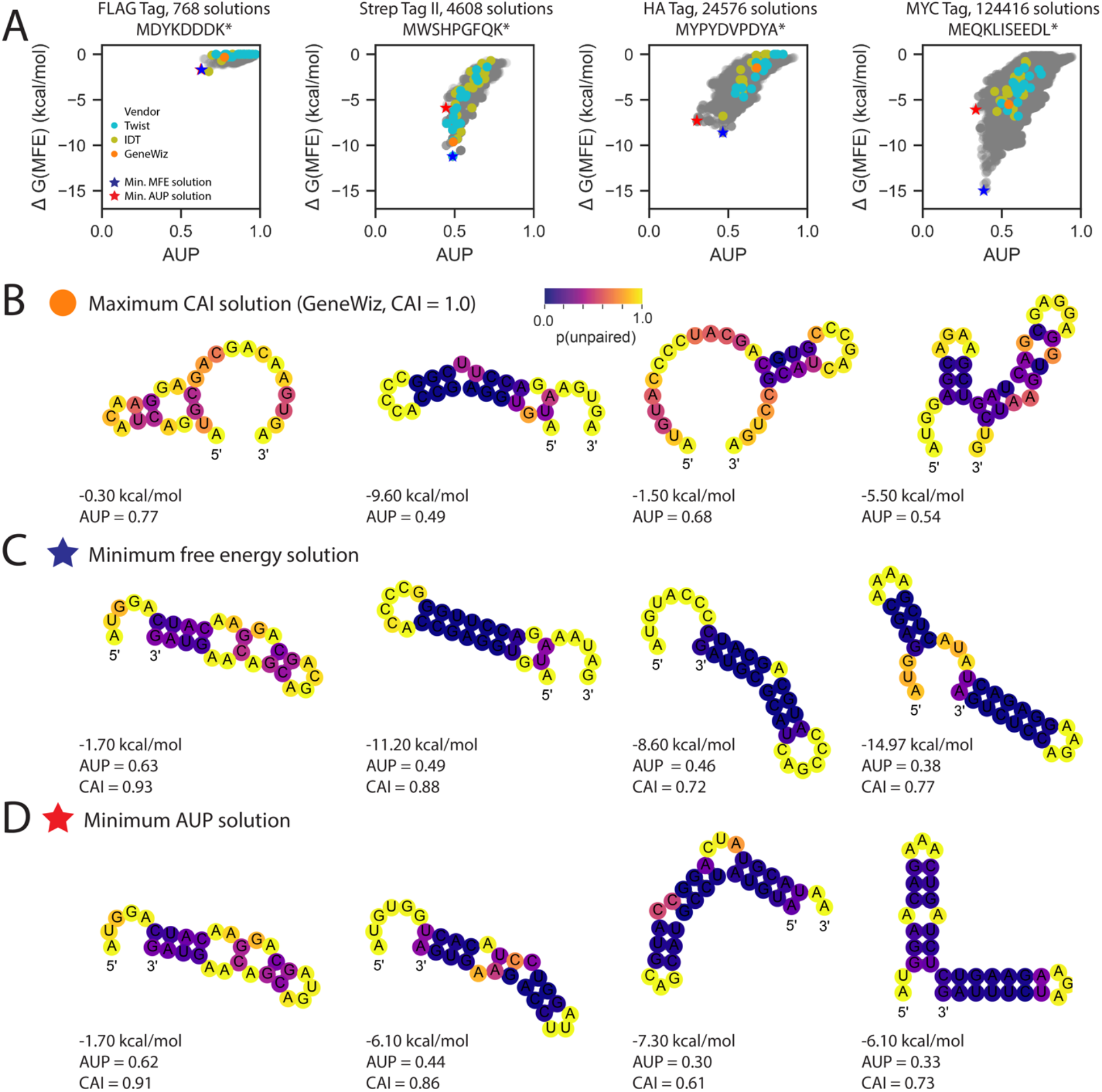
(A) Enumerating all possible coding sequences for small tag proteins reveals that the coding sequence with the lowest energy for its MFE structure (blue star) is not always the same as the coding sequence with the lowest AUP (red star). In (B) – (D), MFE structures for mRNA solutions are shown, with nucleotides are colored by p(unpaired). (B) Sequences from vendors optimizing for codon usage do not reliably minimize ΔG(MFE) or AUP. The mRNA with the lowest-energy MFE structure is displayed in (C), and the mRNA with the lowest AUP is displayed in (D).

**Table 2.**
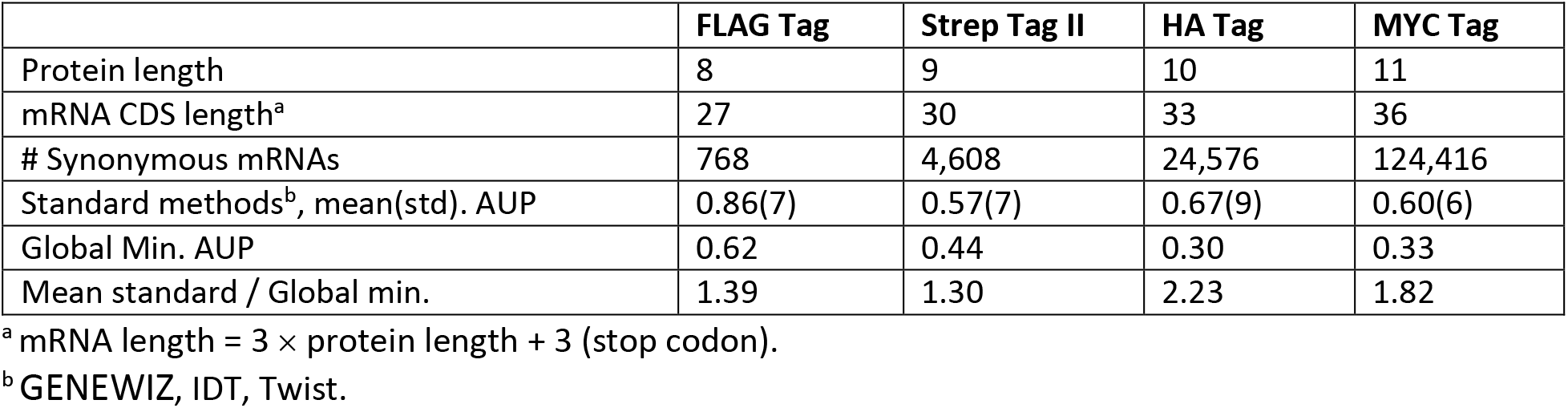
AUP values for example tag proteins and calculated fold-change decrease in AUP from standard design methods to the global minimum AUP value.

|  | **FLAG Tag** | **Strep Tag II** | **HA Tag** | **MYC Tag** |
| --- | --- | --- | --- | --- |
| Protein length | 8 | 9 | 10 | 11 |
| mRNA CDS length<sup>a</sup> | 27 | 30 | 33 | 36 |
| # Synonymous mRNAs | 768 | 4,608 | 24,576 | 124,416 |
| Standard methods<sup>b</sup>, mean(std). AUP | 0.86(7) | 0.57(7) | 0.67(9) | 0.60(6) |
| Global Min. AUP | 0.62 | 0.44 | 0.30 | 0.33 |
| Mean standard / Global min. | 1.39 | 1.30 | 2.23 | 1.82 |

<sup>a</sup> mRNA length = 3 × protein length + 3 (stop codon).
<sup>b</sup> GENEWIZ, IDT, Twist.

Interestingly, we discovered that minimal-AUP solutions were close to optimal with respect to other metrics previously considered in mRNA design. The codon adaptation index (CAI), or related metrics that match codons frequency of the designed mRNA to mRNA’s in the target organisms, are optimized by vendor design algorithms (33); the GENEWIZ algorithm gives a CAI of 1.0. The minimum AUP solutions have CAI values of 0.91, 0.86, 0.61, and 0.73 respectively, which are comparable to vendor-generated sequences, suggesting that optimization of AUP will not result in a major penalty on CAI.

In other studies, structured mRNA’s have been designed through optimization of the predicted folding free energy of the minimum free energy structure (MFE), which can be achieved through an exact algorithm (28–30). For 4 of the 10 model systems studied – the three His tags and the FLAG tag – the coding sequence with the lowest free energy MFE structure also exhibited the lowest AUP (Figure 2A, Figure S1). For other model proteins, including the Strep-tag II, the HA tag, and the Myc tag, the coding sequence with the lowest-energy MFE structure was not the same as the solution with the lowest AUP (Figure 2A). The most notable difference was in designs for the HA tag: the lowest AUP value obtained was 0.30, which is 1.5-fold lower than AUP of the minimal MFE solution, 0.46. However, inspection of the two solutions clarifies why a structure with a higher free energy but a lower AUP would be preferred if we wish to reduce overall hydrolysis (Figure 2C, 2D). Minimal AUP solutions have more stems and fewer ‘hot spots’ (7 vs. 15 yellow nucleotides in HA panel of Figures 2D vs. 2C) rather than optimize the folding free energy of each stem, once formed (reflected here in the base pairing probability; magenta vs. dark purple coloring in HA panel of Figures 2D vs. 2C). Repeating the same analysis across other small model systems as well as with other secondary-structure packages (27, 34) reveals similar results (Figure S1). Taken together, this enumerative analysis of small model mRNAs suggested up to 2-fold increased stabilization might be achievable in mRNA design while retaining excellent codon adaptation indices and that solutions with minimal folding free energy are not necessarily expected to be most stable to hydrolysis.

### A two-fold decrease in AUP is achievable for long mRNA constructs

To test the applicability of our insights from small peptide-encoding mRNA’s to more realistic protein-encoding mRNA design problems, we tested mRNA’s with lengths of hundreds of nucleotides encoding a variety of target proteins, some with therapeutic potential against SARS-CoV-2 and some commonly used in laboratory settings and animal studies to test protein synthesis levels (Figure 1C). The four systems were a multi-epitope vaccine design (MEV) derived from SARS-CoV-2 spike glycoprotein (S) and nucleocapsid (N) proteins; Nanoluciferase; enhanced green fluorescence protein with an attached degron sequence (eGFP+deg), used by Mauger et al. (35) for characterizing mRNA stability and translation; and the SARS-CoV-2 spike receptor binding domain (RBD) of the SARS-CoV-2 spike protein. The protein targets of the mRNA design challenges are further described in Methods, and sequences are listed in Table S2. Because enumeration of mRNA sequences is not possible for these problems, we compared sequences generated by a variety of methods (Figure 1D): uniform sampling of codons (“Uniform random codons”); uniform sampling of GC-rich codons only (“GC-rich codons”); vendor-supplied servers from IDT, GENEWIZ, and Twist; the algorithm CDSfold (28), which returns a sequence with minimal ΔG(MFE) solution; the algorithm LinearDesign (29), which returns a minimal ΔG(MFE) solution that is weighted by codon optimality, as well as sequences from other groups when possible (35). The algorithm CDSfold has the option to alter the maximum allowed base pair distance; we varied this parameter for each design challenge to test if AUP from CDSfold designs varied with maximal allowed base pair distance. We further developed a stochastic Monte Carlo Tree Search algorithm, RiboTree, to stochastically minimize AUP of model mRNAs (see Methods). Last, we crowdsourced solutions through the online RNA design platform Eterna (36). Early Eterna challenges, labeled “Eterna, exploratory” in Figures 3 and 4, were not set up with any specific optimization targets other than a general call to create mRNAs that coded for the target proteins but formed significant structures in Eterna’s game interface, which provides folding calculations in a number of secondary structure prediction packages (see Methods). An additional set of Eterna sequences were solicited in the “p(unp) challenges”, where the AUP metric was calculated and provided to Eterna participants within the game interface to guide optimization.

**Figure 3.**
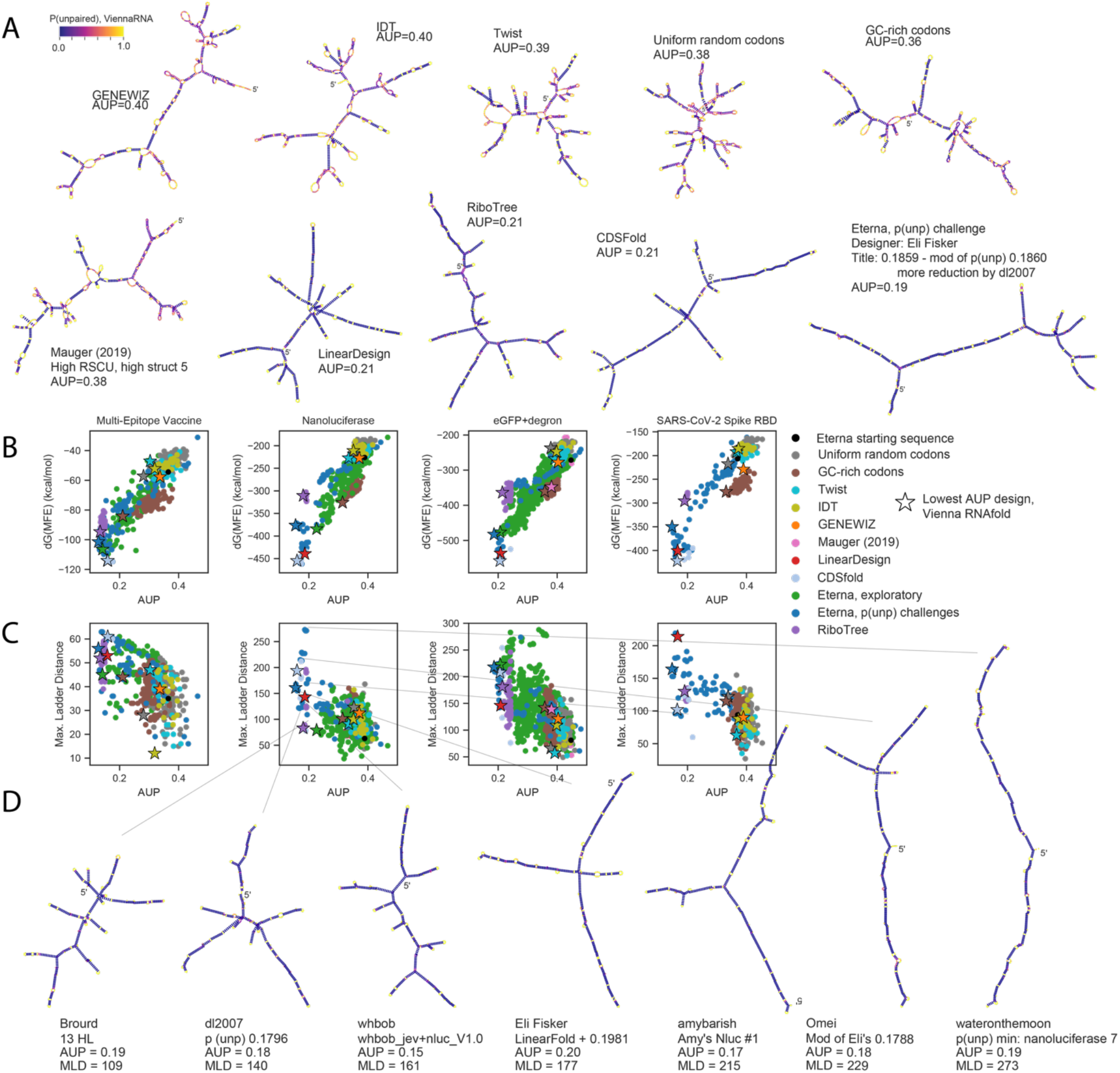
Sequences designed rationally by participants during Eterna’s OpenVaccine challenge result in the lowest AUP values for mRNAs encoding a variety of model proteins used for studying translation and as model vaccines, ranging in length from 144 nucleotides (the Multi-epitope Vaccine) to 855 nucleotides (eGFP+degron (35)). (A) Force-directed graph visualization of sequences coding for eGFP+degron with lowest AUP value from each design source, colored by AUP per nucleotide. (B) While ΔG(MFE) and AUP are correlated, the design with the lowest AUP is not the same as the design with the lowest ΔG(MFE). Starred points indicate the design for each design strategy with the lowest AUP value, calculated with ViennaRNA. (C) Eterna designs show structural diversity as characterized by the Maximum Ladder Distance (MLD), the longest path of contiguous helices present in the minimum free energy (MFE) structure of the molecule. (D) MFE structures predicted in the ViennaRNA structure prediction package are depicted for designs a variety of MLD values, indicating similarly stabilized stems for a range of topologies.

**Figure 4.**
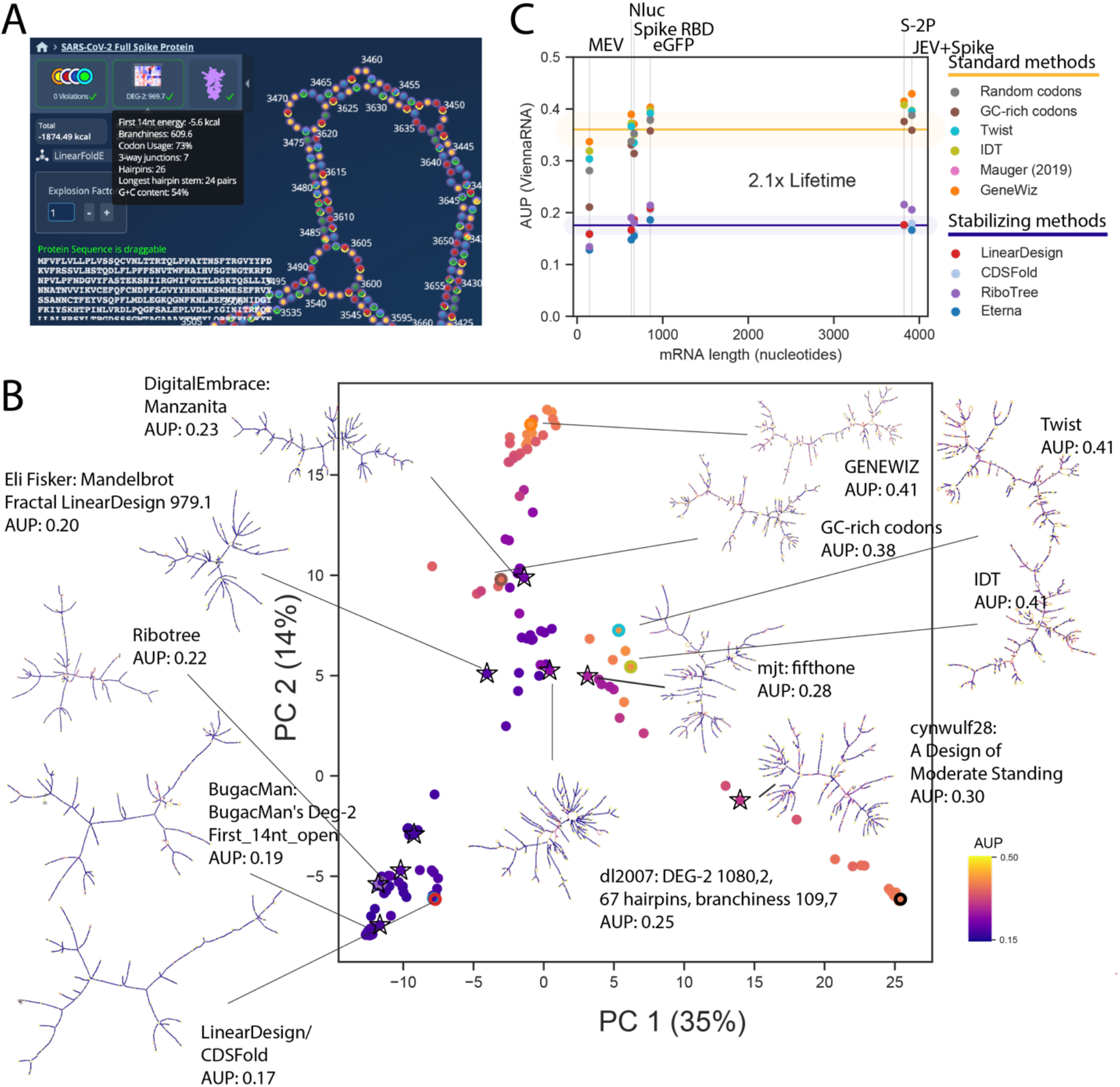
Design of stabilized mRNAs for the SARS-CoV-2 full spike protein achieve the same degree of stabilization as in smaller mRNA design challenges. (A) Eterna participants were provided with metrics on a variety of structure metrics described in this work to aid in creating designs with a diversity of values. (B) Solutions voted upon by the Eterna community show a diversity of structures while maintaining low AUP values. The solutions with the lowest AUP are structurally similar to and were derived from the ΔG(MFE) optimal structure from LinearDesign. (C) AUP values from different design methods are consistent across different mRNA lengths. A two-fold increase in lifetime is predicted by changing from a “Standard” design method (methods that do not stabilize structure) to a design method that increases structure.

For all four challenges, the sequences with the lowest AUP values were designed by Eterna participants. We found that designs from the tested algorithmic and crowdsourcing approaches encompassed a wide range of sequence space, and that sequences with low AUP values did not localize to specific regions of sequence space (Figure S2). Figure 3A depicts MFE structures of the minimal AUP sequence for each design method for the eGFP+degron challenge (the longest mRNA), with nucleotides colored by their unpaired probability, as calculated in the ViennaRNA folding package (24). MFE structures of minimal AUP sequences from each mRNA challenge are in Figure S3. Structures portrayed in Figure 3A indicate visual hallmarks of structures with lower AUP: solutions from LinearDesign, CDSFold, RiboTree, and Eterna have longer helices, fewer loops and junctions, and lower unpaired probabilities in stems (indicated by dark purple). Notably, the solutions with the minimal AUP were distinct from solutions with the lowest ΔG(MFE) (Figure 3B) for all four challenges. Table 3 contains summary statistics for AUP values for design methods separated by standard methods (codon sampling, gene vendor tools) and methods intended to stabilize secondary structure (Eterna p(unp) rational design, CDSfold, LinearDesign, RiboTree).

**Table 3.** Statistics of AUP values obtained in comparing different classes of design methods on mRNA design challenges in this study.

| mRNA design challenge | Multi-epitope vaccine | Nano-luciferase | eGFP + degron | Spike RBD | JEV + Spike | S-2P |
| --- | --- | --- | --- | --- | --- | --- |
| Protein length (aa) | 47 | 221 | 284 | 210 | 1303 | 1273 |
| mRNA CDS length (nt) | 144 | 666 | 855 | 633 | 3912 | 3822 |
| Standard methods<sup>b</sup>, mean(std) AUP | 0.34(4) | 0.36(2) | 0.40(2) | 0.39(2) | 0.40(2) | 0.40(2) |
| Stabilizing methods<sup>c</sup>, mean(std) AUP | 0.15(2) | 0.17(2) | 0.20(1) | 0.17(2) | 0.18(2) | 0.19(2) |
| Stabilizing methods<sup>c</sup>, min. AUP<sup>d</sup> | 0.13 | 0.15 | 0.19 | 0.15 | 0.17 | 0.17 |
| Mean standard / min stabilized | 2.6 | 2.4 | 2.1 | 2.6 | 2.4 | 2.4 |

<sup>a</sup> mRNA length = $3 \times$ protein length + 3 (stop codon).
<sup>b</sup> Uniform codons, GC codons, GENEWIZ, IDT, Twist.
<sup>c</sup> Eterna p(unp) challenge, RiboTree, LinearDesign.
<sup>d</sup> In all cases, minimal AUP was achieved in Eterna p(unp) challenge.

The values of AUP achieved by Eterna participant submissions in the “p(unp) challenge” (mean and standard deviations of MEV: 0.22 ± 0.08, Nluc: 0.24 ± 0.08, eGFP: 0.28 ± 0.08, Spike RBD: 0.24 ± 0.08) were significantly lower than values from standard methods, including codon random sampling and vendor-generated sequences (MEV: 0.34 ± 0.04, Nluc: 0.36 ± 0.02, eGFP: 0.40 ± 0.02, Spike RBD: 0.39 ± 0.02, Table 3). The lowest AUP values from Eterna participants (MEV: 0.128, Nluc: 0.155, eGFP: 0.186, Spike RBD: 0.148) were lower in each case than the AUP values of LinearDesign constructs, (MEV: 0.159, Nluc: 0.186, eGFP: 0.208, Spike RBD: 0.167), CDSfold constructs (MEV: 0.160, Nluc: 0.160, eGFP: 0.206, Spike RBD: 0.165), or of minimum AUP solutions from RiboTree (MEV: 0.134, Nluc: 0.181, eGFP: 0.214, Spike RBD: 0.190). RiboTree came closest (within 5%) to the minimal Eterna AUP value for the shortest mRNA sequence, suggesting that RiboTree was better able to search sequence space for the shorter sequences.

One of the challenges, the eGFP+degron mRNA, could be compared to designs developed by Mauger et al. based on folding free energy optimization to increase functional mRNA lifetime in human cells (35). The minimal AUP value from those sequences (0.381) was similar to the value obtained from randomly-sampled codons, indicating that explicit optimization of AUP rather than folding free energy is necessary for applications seeking stability against hydrolysis. Repeating these analyses of mRNAs with other secondary structure packages (27, 34) reveals similar results (Figure S4).

We were interested to note that RiboTree solutions exhibited low AUP while not necessarily minimizing ΔG(MFE). Minimum AUP solutions from RiboTree had absolute ΔG(MFE) values that were up to 75% reduced (less stable) compared to absolute ΔG(MFE) values of minimum ΔG(MFE) solutions, which came from Eterna participants (MEV: 93%, Nanoluciferase: 82%, eGFP+deg: 75%, Spike RBD: 84%). Minimizing AUP without minimizing ΔG(MFE) may prove to be a valuable design strategy for developing mRNAs that are stable under storage but need to be sufficiently unstable as to exhibit cooperative unfolding by the cells’ translational apparatus.

### Diversity of properties related to translation and immunogenic function

After establishing the feasibility of designing mRNA sequences with reduced AUP, we wished to determine if these sequences might be viable for translation and for either preventing or eliciting innate immune responses. In advance of experimental tests, we tabulated sequence and structure properties that have been hypothesized to correlate with translation and immunogenicity.

We first characterized the codon adaptation index (CAI) (33) of sequences across design methods, as this measure has been implicated in improving translation efficiency. We found that across all p(unp) challenges, minimal AUP sequences consistently had CAI values greater than 0.7 (Figure S5). Another design feature that has been hypothesized to influence protein translation efficiency is the exposure of the CDS immediately upstream of the initiation codon (37, 38). We calculated the average unpaired probabilities of the first 14 nucleotides (37), termed AUP^init,14^, in the presence of our model UTRs (HBB). A higher value of AUP^init,14^ indicates a more exposed ribosome initiation site, and is expected to correlate with higher translation efficiency. We found a range of AUP^init,14^ values possible for low AUP sequences (Figure S5). These analyses suggest that it is feasible to design low AUP sequences that are translatable, as assessed by the available metrics of CAI and AUP^init,14^.

Another important consideration is the possibility of mRNA therapeutics eliciting immunogenic responses from pathways that recognize double-stranded RNA helices. We found that none of the sequences characterized included helices longer than 33 base pairs, a measure that has been found to be the minimum length that leads to global shutdown of cellular mRNA translation after sensing by protein kinase R (39) (Figure S5). Importantly, however, different low-AUP designs exhibited a wide spectrum of helix lengths (8 to 22 base pairs) suggesting that a less drastic innate immune response might be achieved and the response may be tunable depending on whether such responses are desirable (mRNA vaccines) or not (e.g., for anti-immune mRNA therapeutics).

Finally, the sequences designed in the above challenges did not contain untranslated regions (UTRs). We compared the AUP of the above designs in the presence and of standard human beta globin (HBB) UTRs, as well as AUP^init,14^, as the presence of a 5’ UTR could base pair with a ribosome binding site. We found that for the collected sequence designs, the AUP calculated in the context of HBB UTRs had high correlation to the AUP of the CDS only (MEV: 0.91, Nanoluciferase: 0.98, eGFP+degron: 0.99, Spike RBD: 0.99, Figure S6, UTR sequences in Table S2). This indicates that the overall AUP of a designed coding sequence (CDS) also maintains low AUP in the context of UTRs. The correlation between AUP^init,14^ in the absence and in the presence of UTRs was less robust (MEV: 0.32, Nanoluciferase: 0.57, eGFP+degron: 0.56, Spike RBD: 0.71, Figure S6), but still suggests that constructs may be designed to maintain high AUP^init,14^ that is robust to adding UTRs.

In addition to structural characteristics that affect stability against in vitro hydrolysis, translatability and degradation rates in cells, and immunogenicity of mRNA molecules, we expect there are many structural characteristics that relate to a molecule’s in vivo persistence that are not yet well understood. The ability to design multiple low-AUP sequences with a large range of alternative structures increases the potential that a functional design may be found in empirical tests or as the connections between mRNA structure and function are better understood. For instance, in Figure 3A, we observed that although LinearDesign, RiboTree, and Eterna sequences for an eGFP+degron mRNA all have AUP values within 10% of each other, they have different secondary structures. The same can be seen for all the mRNA design problems we tested (Figure S2).

As a more quantitative evaluation of structural diversity, we characterized the Maximum Ladder Distance (MLD) of designed sequences. This measure has been used to describe the compactness of viral genomic RNAs and has been hypothesized to be relevant for viral packaging, immunogenicity, and biological persistence (39–42). If an RNA molecule’s secondary structure is represented as an undirected graph, where edges represent helices, edge lengths correspond to helix lengths, and vertices correspond to loops, the MLD is the longest path that can be traced in the graph. Genomic viral RNAs have been demonstrated to have shorter MLDs than equivalent random controls, and molecules with shorter MLDs have been shown to be more compact experimentally, a feature that may also contribute to persistence (41). We found that AUP and MLD were negatively correlated across the MEV, Nanoluciferase, eGFP+degron, and Spike RBD challenges (−0.64, −0.59, −0.62, −0.70, respectively, Figure 3C). This overall (negative) correlation reflects how minimizing AUP leads to larger average MLD values. Nevertheless, we note that the MLD values still fall over a wide range for sequences with low AUP. Example structures from the Nanoluciferase challenge, depicted in Figure 3D, range from highly branched, compact structures (Figure 3D, left) to long, snake-like structures (Figure 3D, right). These structures exhibit uniformly low unpaired probabilities in stems (indicated by dark purple coloring), with the main difference being the layout of stems. In addition to MLD, we calculated several other metrics characterizing structure, such as counts of different types of loops and junctions, the ratio of number of hairpins to number of 3-way junctions in the MFE structure, introduced in ref. (41) as a measure of branching, and mean distance between nucleotides in base pairs. In all cases, values ranged by over 2-fold in low-AUP solutions, underscoring the diversity of structures that can be achieved. Statistics of these metrics across the mRNA challenges and different design methods are provided in Figure S7.

These results demonstrate that both automated and rational design methods are capable of finding RNA sequences with low AUP values but a wide range of diverse structures. Testing these mRNAs experimentally for their translation rates and persistence in cells and in animals will help address the relationship between MLD and mRNA therapeutic stability.

### Eterna participants are able to design stabilized SARS-CoV-2 full spike Protein mRNAs

For longer mRNA design problems, including the SARS-CoV-2 spike protein mRNA used in COVID-19 vaccine formulations (3822 nts), we noted that the computational cost associated with computing thermodynamic ensembles associated with AUP became slow and hindered automated or interactive design guided by AUP. We therefore sought other observables that were more rapid to compute to guide design of RNA’s stabilized against hydrolysis. We calculated correlations between many observables and AUP (Figure S8), and found that for all four challenges, the number of unpaired nucleotides in the single MFE structure was the most correlated with AUP, giving near-perfect correlations (0.98, 0.99, 0.99, 0.99, respectively). We leveraged this observation to launch another design puzzle on Eterna: minimizing the number of unpaired nucleotides in the MFE structure, as a proxy for AUP, for a vaccine design that includes the full SARS-CoV-2 spike protein (“JEV + Spike”). We found that Eterna participants were capable of finding values for AUP as low as in previous challenges, despite the fact that the JEV + Spike mRNA was over four times as long as previous challenges. Again, this solution was distinct from the minimal ΔG(MFE) design calculated in CDSFold (Figure S9). The lowest AUP value for the JEV + Spike protein was 0.166, 2.5-fold lower than the minimum AUP values from conventional design methods (0.40 +/- 0.02). RiboTree was also run minimizing the number of unpaired nucleotides. The larger size of the JEV+Spike protein meant that it took longer for RiboTree to minimize the solution. Starting from a random initialization and running RiboTree for 6000 iterations (2 days) resulted in a construct with an AUP of 0.254. Seeding RiboTree with a starting sequence partially stabilized using the LinearDesign server (AUP: 0.212) resulted in reducing the AUP (0.206).

The SARS-CoV-2 spike protein sequence used in most vaccine formulations has turned out to be a double proline mutant S-2P that stabilizes the prefusion conformation of the 1273-aa spike (43). We launched one more puzzle on Eterna calling for solutions for stabilized mRNAs encoding the S-2P protein. Participants were provided with a variety of metrics of both predicted stability and structure (Figure 4A, Methods), and were not specifically asked to optimize any of the metrics. Out of 181 submissions, the top 9 solutions that were voted upon demonstrated a diverse set of sequences, some prioritizing structure diversity, some prioritizing high stability, all demonstrating low AUP values (Figure 4B). As with shorter mRNAs, S-2P solutions with the lowest AUP values – from Eterna participants, RiboTree, CDSfold, and LinearDesign – demonstrate a 2-fold reduction in AUP from mRNAs designed through randomly selecting codons or from codon optimization algorithms from gene synthesis vendors (Figure 4C).

### Highly stable mRNAs are robust to variations in design

There are several design contexts in which it would be advantageous to adapt an existing highly stable mRNA design, rather than design an mRNA *de novo*. These include changes in environment (e.g., higher temperature), changes to protein sequence, potentially needed to rapidly develop booster vaccines for variant strains (44), as well as altering the UTR used, which would allow for flexibility in testing different expression formulations post-mRNA-design. We tested the robustness of designed S-2P mRNA stability to small changes in protein sequence and to different UTRs for a subset of the S-2P sequences collected. We selected the 9 top-voted sequences from the Eterna S-2P round, as well as one representative sequence from other methods (Twist, IDT, GENEWIZ, GC-rich, LinearDesign, and RiboTree).

As a test of design robustness to destabilizing environments, we compared the predicted AUP at the default temperature of our folding packages(37 °C) to the predicted AUP at 70°C. Sequences with low AUPs maintained low AUP predicted at 70°C, while sequences with high AUPs had AUPs raised by roughly an additional 30% (Figure S10).

To test robustness of folding stability with respect to nonsynonymous coding changes, we tested a simple heuristic to ‘hot fix’ an mRNA design to code for a new protein variant: for each amino acid mutation, we replaced the new codon with the most GC-rich codons for the mutant. We used this heuristic to design mRNA sequences coding for S-2P antigens appropriate strains B.1.1.7, (45) P.1.(46), and B.1.351 (47), which present 10, 10, and 12 amino acid changes compared to S-2P, respectively (Protein sequences listed in Table S2). We found that for the mRNAs tested, the AUP of the modified mRNA had near-perfect correlation with the AUP of the original mRNA (0.999, 0.996, 0.999, for B.1.1.7, P.1, B.1.351, respectively). The addition of mutations did not perturb the global layout of the mRNA design (example for one sequence depicted in Figure 5, and a second in Figure S11). Unlike in the temperature test, less-structured sequences also retained very similar AUP values. To test the effect of adding UTRs, we calculated AUP for the subset of sequences in the context of human hemoglobin subunit beta (HBB) 5’/3’ UTR. The AUP of the full constructs exhibited very high correlation to the AUP of the CDS only (0.999 respectively). Again, less-structured sequences also retained very similar AUP values. Taken together, these results indicate that for messenger RNAs of similar length to the SARS-CoV-2 spike protein (3822 nts), predicted increases in stability are robust to protein sequence mutations and changes in UTRs.

**Figure 5.**
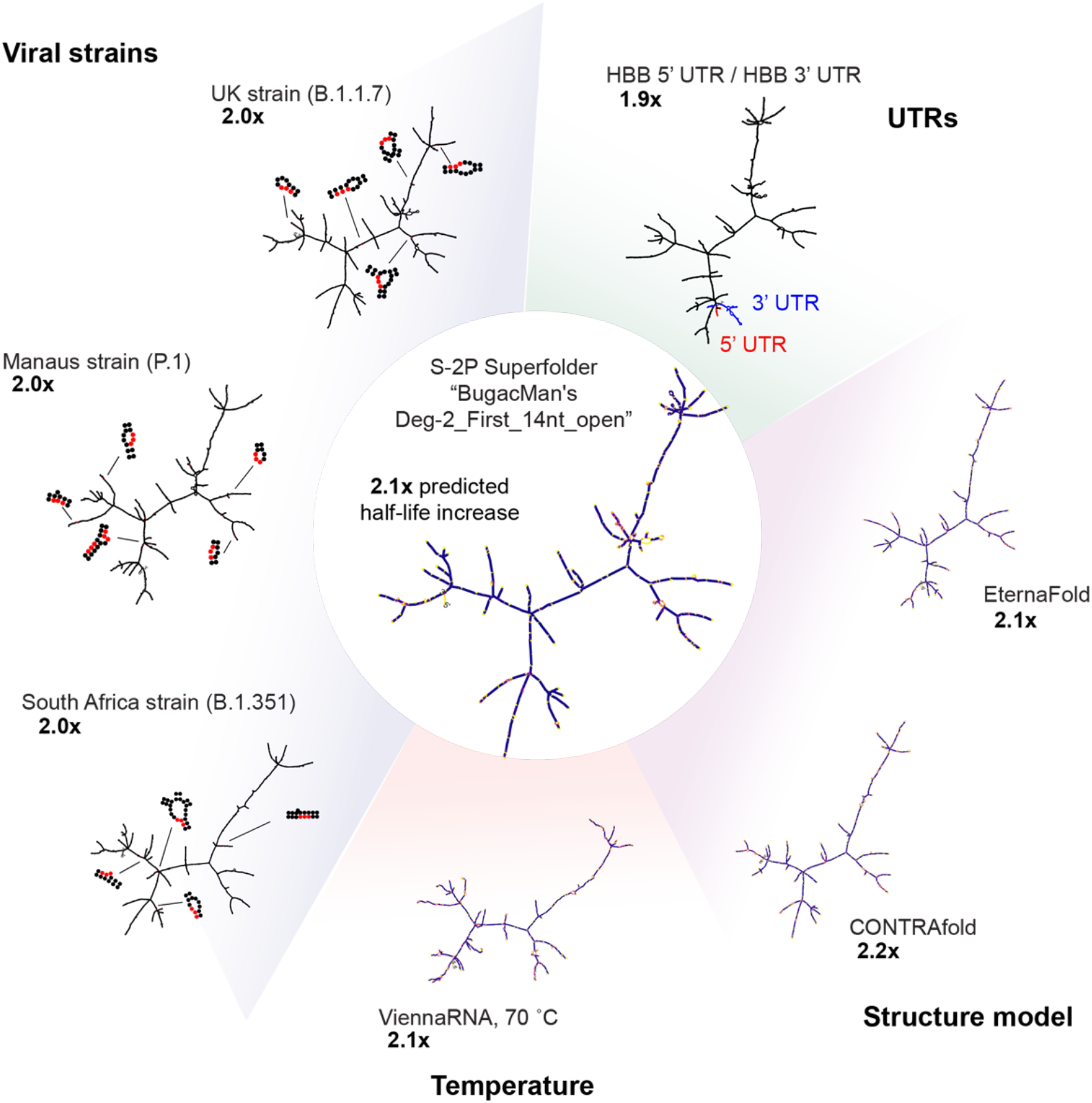
Low AUP solutions as superfolder mRNAs. Stabilization derived from a low AUP solution is robust to small variations in protein sequence, added untranslated regions (UTRs), choice of folding algorithm, and higher temperatures.

## Discussion

In this work, we have developed a complete framework for stabilizing messenger RNA against in vitro hydrolysis through structure-based design. We presented a model relating the degradation rate of an RNA molecule to the average of unpaired probabilities (AUP), a measure readily calculated in available secondary structure prediction packages. We calculated AUP across a large collection of messenger RNA designs for small peptides, for reporter proteins like eGFP, and for antigens under consideration for SARS-CoV-2 vaccines. The solutions with the lowest AUP values – from Eterna participants, the new RiboTree algorithm, and LinearDesign – demonstrate a 2-fold reduction in AUP from mRNAs designed through randomly selecting codons or from codon optimization algorithms from gene synthesis vendors (Figure 4C). A 2-fold reduction in AUP corresponds to a 2-fold increase in mRNA half-life, a potentially significant improvement in the context of mRNA biotechnology and current logistical challenges facing mRNA vaccine distribution for the COVID pandemic: extra days or weeks of stability of COVID mRNA vaccines in aqueous buffer could dramatically increase the number of people who can receive doses and potentially obviate the need to ship the vaccines in frozen vials (8).

Important for these practical applications, mRNAs stabilized through designed secondary structure must remain functional. The translational efficiency and immunogenicity of mRNA molecules remain difficult to predict from sequence and structure, so it is important that when designing for one property (i.e., low hydrolysis, as investigated in this work), a wide range of values for other sequence and structure features should be achievable. We were encouraged to find that low AUP designs do indeed encompass a variety of structures, as measured by maximum helix length, maximum ladder distance, number of multiloop junctions, and numerous other properties. Importantly, if mRNA design efforts require maximizing or minimizing these sequence or structural metrics, e.g., to enhance packaging into lipid nanoparticles or to suppress innate immune responses, both Eterna crowdsourcing, dynamic programming approaches such as CDSFold and LinearDesign, and the automated RiboTree framework allow for optimization of such properties simultaneously with AUP. Finally, we demonstrate that for mRNA designs at the length scale of the SARS-CoV-2 spike protein, stabilized mRNA designs are robust to a number of changes: increasing the temperature, altering the protein sequence (potentially useful for developing ‘booster’ vaccines for variant strains), and adding different UTR sequences to the designed CDS. By analogy to ‘superfolder’ proteins that are stabilized against similar perturbations of environment and sequence, we propose to call mRNA sequences designed to have the same properties ‘superfolder’ mRNAs.

Further increases in mRNA lifetime through structure-guided design are likely possible, as the computational model underlying our study is expected to underestimate how much stabilization is achievable in practice. In particular, some secondary structure and sequence motifs may be less prone to hydrolysis than others (11, 16, 22). Knowledge and prioritization of those specially hydrolysis-resistant motifs in mRNA designs could lower degradation rates beyond those achieved in the present study. Critical for achieving such improvements in stability in practice will be collection of large experimental data sets mapping hydrolysis rates of many RNA’s at single nucleotide resolution, and predictive models of hydrolysis rates trained on such data sets. Measurements of protein expression from superfolder mRNAs will also be important to test compatibility with cellular translation and sustained or increased cellular lifetime, as has been recently observed (35). We propose that with such empirical knowledge, mRNA lifetimes in storage and shipping may be extended by much more than two-fold with maintained or enhanced function.

## Dedication

This paper is in memory of Malcolm Watson, one of Eterna’s most active participants and longtime contributors.

## Acknowledgements

We thank Jonathan Romano, Sharif Ezzat, and Camilla Kao for Eterna development and assistance launching the OpenVaccine challenge and the AUP metric on the Eterna platform. We thank Ivan Zheludev for advice in designing the SARS-CoV-2 multi-epitope vaccine protein. We thank Eesha Sharma, Wipapat Kladwang, Kathrin Leppek, Gun Woo Byeon, Craig Kerr, Daphna Rothschild Bup, other members of the Das and Barna labs (Stanford), and Mike Jewett (Northwestern University) for useful discussion regarding mRNA design challenges, degradation and translation. We thank Goro Terai and Kiyoshi Asai (National Institute of Advanced Industrial Science and Technology (AIST), Koto-ku, Tokyo) for discussions of CDSFold. We thank Liang Huang (Oregon State University, Baidu Research USA) for discussions of LinearDesign. RiboTree calculations were performed on the Stanford Sherlock cluster. We acknowledge funding from the National Science Foundation (GRFP to H.K.W.S. and D.S.K.), Stanford University Graduate Research Fellowship (C.A.C.), the National Institute of Health (R35 GM122579 to R.D.), FastGrants, and gifts to the Eterna OpenVaccine project from donors listed in Table S3.

## Contributions

H.K.W.S., D.S.K., C.A.C., R.D. designed and conceptualized the research. D.S.K., A.M.W., R.W.O., and J.J.N. developed the Eterna interface, launched the Eterna puzzles, collected sequences, and solicited feedback from Eterna participants. C.A.C. and D.S.K. developed the RiboTree software and performed RiboTree runs. R.A.P.S. and P.H. designed the protein sequence used in the SARS-CoV-2 Spike receptor binding domain mRNA challenge. H.K.W.S. performed analysis and developed the *OpenVaccine-solves* dataset and metrics. H.K.W.S., D.S.K., and R.D. wrote the manuscript.

## Methods

### Data Availability

The OpenVaccine sequences and calculated features are included in the supplementary information of this manuscript. The same data, as well as scripts to reproduce analysis, are available in the “OpenVaccine-solves” database under an Open COVID license at https://eternagame.org/about/software.

### Quantitative model for RNA hydrolysis

To predict the RNA degradation rates in Table 1, we used a model presented in ref. (14) for an inherent rate for phosphodiester bond cleavage as a function of pH, temperature, and ionic concentrations. The model is reproduced below:

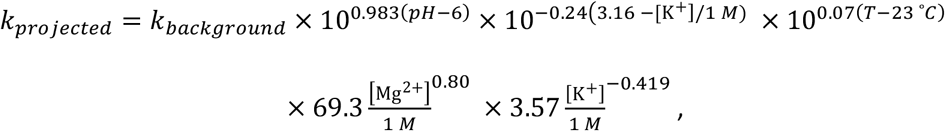

where *k_background_* = 1.30 × 10^−9^ min^−1^, which represents the model’s selected reference point: pH 6, 23 ° C, [K^+^] = 3.17 M. The above equation was parametrized from measurements with [Mg^2+^] concentrations between 0.005 and 0.01 M; for conditions with 0 M Mg^2+^, we omit the terms in the second line.

### Target identification for mRNA design and optimization

Messenger RNAs encoding five target proteins were chosen for this work: 1) a candidate multi-epitope vaccine design (MEV) derived from SARS-CoV-2 Spike and Nucleocapsid proteins, 2) Nanoluciferase, 3) eGFP with a degron sequence (eGFP+deg), 4) the Receptor Binding Domain (RBD) of the SARS-CoV-2 Spike protein, and 5) the full SARS-CoV-2 Spike protein. Here we provide in detail the rationale behind how the constructs were chosen and designed.

Nanoluciferase and eGFP+degron were chosen for their ubiquitous use in biomedical research. The Nanoluciferase sequence was taken from ref. (48). The choice of eGFP+degron allowed for close comparison with designed mRNAs from ref. (35), which investigated the relationships between mRNA structure, stability and translation using the eGFP+degron, while Nanoluciferase was chosen to complement another easily assayed, commonly used model protein in mammalian systems.

For the MEV protein, we chose peptide sequences from SARS-CoV with verified T cell-positive assays that were identical in SARS-CoV-2, and one that maximized population coverage according to MHC allele mapping(49). Two of the epitopes (GYQPYRVVVL and PYRVVVLSF) map directly onto the receptor binding domain of the S (spike) protein, which we hypothesized to be important in eliciting the proper antibody/immune response. The last epitope (LSPRWYFYY), from the N protein, was chosen for its potentially high coverage of effectiveness in the global population. To test our hypothesis against a wide range of mRNA lengths, we chose to include only three peptide epitopes to introduce a shorter mRNA in our MEV design, but it should be noted that a more realistic MEV may feature many more epitopes (50, 51).

The SARS-CoV-2 Spike RBD is derived from the reported structure of the spike protein in the prefusion conformation(52). Similarly, the protein chosen for the “JEV + spike” protein is not the full sequence as found in the genome of SARS-CoV-2(1), but as reported in the pre-fusion conformation, which is hypothesized to promote an enhanced immune response in use as a vaccine. For three of the proteins (Nanoluciferase, RBD, and “JEV+ Spike” protein), the signal sequence from the Japanese encephalitis virus (JEV) was added to the N-terminus to encode designs for a secreted vaccine (53). The “JEV + Spike” protein additionally had StrepII tag appended.

### Eterna puzzle deployment

Eterna puzzles were launched in a series of rounds that gradually increased the complexity of the sequences designed and the information provided, as outlined in Table 4. For all puzzles, the MFE structure calculated in the default folding engine is displayed as participants design the mRNA molecule.

**Table 4.** Eterna OpenVaccine rounds, included mRNA design challenges, default folding engine, and default metrics provided to participants.

| Title | Constructs | Metrics provided | Default folding engine | URL |
| --- | --- | --- | --- | --- |
| [Lightning Round] Eterna takes on the Coronavirus | MEV, eGFP | $\Delta G(\text{MFE})$, $\Delta G(\text{ensemble})$ | ViennaRNA | https://eternagame.org/labs/9818634 |
| [Round 2] Design of eGFP and epitope mRNA molecules | MEV, eGFP | $\Delta G(\text{MFE})$, $\Delta G(\text{ensemble})$ | ViennaRNA | https://eternagame.org/labs/9833861 |
| [Round 3-extended] Lightning Round Design + Vote of Nanoluciferase | eGFP, Nluc | $\Delta G(\text{MFE})$, $\Delta G(\text{ensemble})$ | ViennaRNA | https://eternagame.org/labs/9874218 |
| [Round 4] The 5' UTR Design Lab | UTR design (not discussed here) | $\Delta G(\text{MFE})$, $\Delta G(\text{ensemble})$, # hairpins, longest stem, GC content | ViennaRNA | https://eternagame.org/labs/10027854 |
| [Computational lab] Eterna vs. Computers, p(unp) | MEV, Nluc, eGFP, Spike RBD | $\Delta G(\text{MFE})$, $\Delta G(\text{ensemble})$, AUP | LinearFold/LinearPartition (ViennaRNA) | https://eternagame.org/labs/10097602 |
| [Special Design Challenge] Humans vs. Computers take on the full spike protein | JEV+S | $\Delta G(\text{MFE})$, $\Delta G(\text{ensemble})$, # unpaired nucleotides | LinearFold/LinearPartition (ViennaRNA) | https://eternagame.org/labs/10126540 |
| Round 7 | S-2P | DegScore, DegScore Leaderboard, "branchiness", Codon Usage, 3-way junctions, # hairpins, longest hairpin stem, GC content, energy of first 14 nts | LinearFold/LinearPartition (EternaFold) | https://eternagame.org/labs/10506533 |

### Generation of vendor-optimized sequences

The protein sequences for each target (Table S2) were used to generate DNA sequences at Integrated DNA Technologies (IDT, https://www.idtdna.com/CodonOpt), Twist Biosciences (https://ecommerce.twistdna.com/app), and GENEWIZ (https://clims4.genewiz.com/Toolbox/CodonOptimization). For IDT and Twist Biosciences, multiple possible sequences were generated for a given protein sequence, while the GENEWIZ design tool returned one possible optimized DNA/RNA sequence per protein sequence.

### Stochastic minimization of AUP in RiboTree

A Monte-Carlo Tree Search algorithm, named RiboTree, was developed to stochastically minimize AUP for mRNA sequences. RiboTree uses the Upper Confidence bounds applied to Trees (UCT) algorithm(54). The UCT loss function, as applied to the problem of sampling RNA sequences, is

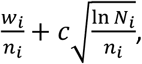

Where *w_i_* is the total score considered for the node after the *i*th move, *n_i_* is the total number of times the node was visited after the *i*th move, and c is the exploration parameter, which determines the tradeoff between depth and breadth search. For minimizing AUP, moves consist of swapping synonymous codons and are accepted with a probability

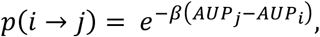

Where *AUP_i_* and *AUP_j_* are the AUP values of states *i* and *j*, respectively, and *β* is a temperature parameter to control the acceptance rate. Runs were terminated after 6000 iterations. The RiboTree code is available for noncommercial use at https://eternagame.org/about/software.

### CDSFold

Solutions from CDSfold were obtained by running the CDSfold algorithm source code with varying maximum base-pairing distance.

### LinearDesign

Solutions from LinearDesign were obtained using the LinearDesign server (http://rna.baidu.com/) and inputting the protein sequences given in Table S2. A maximum beam size of 50 was used for prediction and the standard (Human) codon table.

### Metric calculations

Structure prediction and ensemble-based calculations were performed using LinearFold and LinearPartition with ViennaRNA, CONTRAfold, and EternaFold parameters. Secondary structure features were calculated from predicted MFE structures using RiboGraphViz (www.github.com/DasLab/RiboGraphViz). CAI (Codon Adaptation Index) was calculated as the geometric mean of the relative usage frequency of codons along the length of the coding region, as described in ref (33):

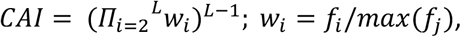

Where *f_j_* represents the frequency of all codons coding for amino acid at position *i*.

### Structure visualization

RNA secondary structures were visualized using draw_RNA (www.github.com/DasLab/draw_rna) and RiboGraphViz (www.github.com/DasLab/RiboGraphViz).

